# Complete telomere-to-telomere assemblies of two sorghum genomes to guide biological discovery

**DOI:** 10.1101/2024.03.24.586445

**Authors:** Chuanzheng Wei, Lei Gao, Ruixue Xiao, Yanbo Wang, Bingru Chen, Wenhui Zou, Jihong Li, Emma Mace, David Jordan, Yongfu Tao

**Author notes:** Correspondence (Yongfu Tao). These authors contribute equally: Chuanzheng Wei, Lei Gao.

## Abstract

Sorghum is a C4 crop well-known for its high efficiency of biomass accumulation and adaptation to drought and hot environments. This study reports the assembly of complete genome sequence of two sorghum genomes using a combination of ultra-long reads from Oxford Nanopore Technology, high-fidelity long reads from Pacbio and Hi-C reads. The accuracy and completeness of the T2T assemblies were comprehensively validated using a range of parameters, including coverage depth assessment, BUSCO, LTR assembly index, ect. Our T2T assembly of BTx623 identified 36.25 Mb of new sequence, in addition to dozens of misoritation and mispositioning of large DNA segments in the BTx623-v3. Correcting these assembly errors in the reference genome is critical for exploiting genetic information in these complex regions in sorghum. Comparison of the two T2T genomes shed new insights on variation of centromeres in sorghum. The two high-quality T2T genomes represent an important resource for sorghum improvement and gene discovery. They would serve as the new reference genomes to unlock the full potential of global sequence variation of this valuable crop.

## INTRODUCTION

Cultivated sorghum (*Sorghum bicolor* L. Moench) is a C_4_ crop well-known for its high efficiency of biomass accumulation and adaptation to drought and hot environments. It is a staple food for half a billion people in Africa and Asia, and provides a major source of feed, fiber and biofuel globally. The release of the first sorghum reference genome of BTx623 greatly accelerated functional genomics studies in sorghum and related C_4_ grasses [1]. Subsequent improvement has further enhanced the quality of the reference genome [2]. Assemblies of other sorghum genomes such as Tx430, Rio and wild sorghum accessions have shown marked intra-specific sequence variation in this crop [3,4]. However, all of the available sorghum genomes are still incomplete, in particular with unresolved centromeres and telomeres, constraining a full understanding of the genomic landscape in the sorghum gene-pool.

## RESULTS

### Assembly of T2T sorghum genomes

In this study, we utilized ultra-long reads from Oxford Nanopore Technology (ONT), high-fidelity (HiFi) long reads from Pacbio, Hi-C reads and Illumina short reads to assemble complete sequences of two sorghum genomes, BTx623 and Ji2055. The white-seeded BTx623 has long served the sorghum community as the reference genome [1], while Ji2055 is an inbred line with red seeds that has led to the successful release of dozens of commercial varieties in China (Figure S1). We generated an average of > 150× sequence coverage of ultra-long ONT data, > 65×coverage of PacBio HiFi data, > 50× HiC data and > 50× Illumina short reads data for both varieties (Table S1). The initial assemblies of the two genomes were obtained using Hifiasm [5] with HiFi reads only, resulting in two genomes containing 246 and 581 contigs, respectively. Hi-C data was then employed to anchor and orient these contigs into 10 pseudomolecules for each genome (Figure S2). Ultra-long ONT reads that were longer than 50 Kb were used together with HiFi reads to fill the sequence gaps and correct assembly errors, which reduced the number of gaps to only four for each assembly. These gaps were then closed via manual extension to achieve gap-free assemblies. Coverage depth analysis using HiFi reads identified 13 genomic regions with assembly errors, which were then corrected using HiFi and ONT reads. After further polishing of the assembled genomes with Illumina reads and HiFi reads, our final telomere-to-telomere (T2T) assemblies were obtained with a genome size of 719.90 Mb for BTx623-T2T (Figure S3) and 722.96 Mb for Ji2055-T2T.

To validate the quality of our T2T assemblies, comprehensive assessments were performed. The overall accuracy of our T2T assemblies was supported by the uniform coverage distribution of PacBio HiFi and ONT reads across nearly all regions of our T2T assemblies (Figure 1A). The two T2T genomes were estimated to have a base accuracy rate of 99.99% using PacBio HiFi reads. The completeness of our T2T assemblies was assessed using the benchmarking universal single-copy orthologs pipeline [6], which showed our two assemblies captured > 98.5% of the 1,614 conserved orthologous genes, slightly higher than the BTx623-v3 (Table S2). LTR assembly index [7], which measures genome continuity, was also higher for our assemblies compared to BTX623-v3 (Table S2). Nearly all the HiFi reads (100%) and ONT (> 99.80%) reads could be mapped back to their derived T2T assemblies, highlighting the completeness of our T2T genomes. The published resequencing data of 44 sorghum varieties [8] was also mapped to the T2T genomes, which displayed a significantly higher mapping rate against our T2T assembles (averaged at 99.20%) than against the BTX623-v3 genome (averaged 97.45%) (Table S3, Figure S4). All the centromeric regions of our T2T genomes contained the sorghum centromere-specific repetitive elements, *PSau3A10* and *pSau3A9* [9, 10] (Figure S5 and Figure S6). Overall, these evidences presented strongly supports the accuracy and completeness of our T2T assemblies.

**Figure 1.**
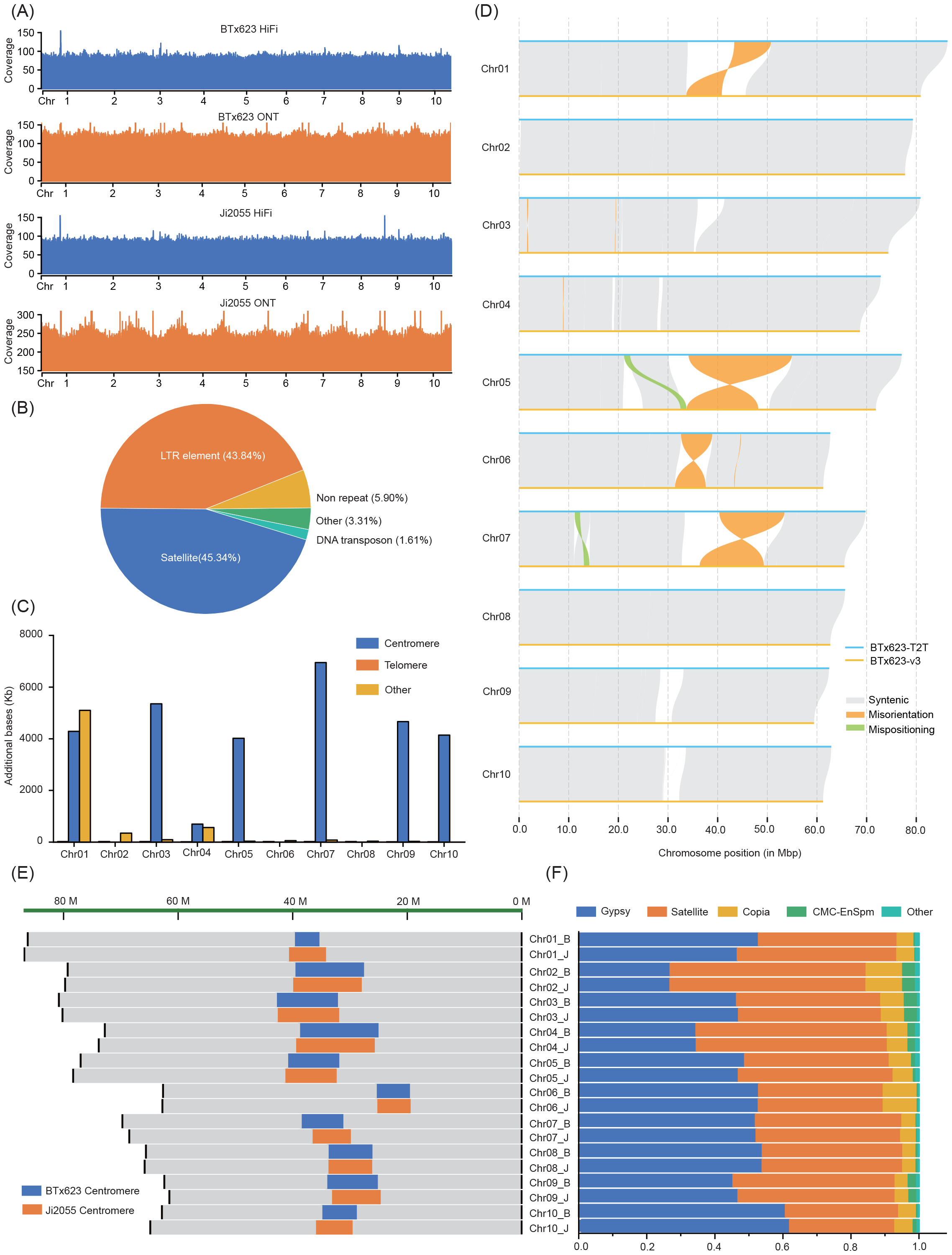
The improvements in sorghum telomere-to-telomere (T2T) genome and variation of centromeres (A) Distribution of coverage depth of high-fidelity (HiFi) reads and Oxford Nanopore Technology (ONT) reads supported the quality of the assembled genomes. (B) Sequence composition of newly identified sequence in BTx623-T2T. (C) Distribution of newly identified sequence in BTx623-T2T. (D) Assembly errors identified in BTx623-v3 according to BTx623-T2T. Only assembly errors larger than 50 Kb were visualized. (E) Variation of centromeres between BTx623-T2T and Ji2055-T2T. (F) Sequence composition of centromere sequence in BTx623-T2T and Ji2055-T2T. The chromosome names contained two parts. The part before the dash sign is chromosome number. The letter after the dash sign represents the genome. Letter “B” represents BTx623-T2T genome. Letter “J” represents Ji2055-T2T genome.

These two T2T sorghum assemblies with their complete genome sequence and intact centromeres and telomeres of all 10 chromosomes represent a significant improvement over the previous version of the reference genome (Table S4 and S5). Genome annotation showed repetitive elements accounted for 66.50% of the BTx623-T2T genome and 65.22% of the Ji2055-T2T genome, including around 50% retroelements and 9% DNA transposons (Table S6). Our T2T assemblies contained slightly higher percentage of repetitive elements than BTx623-v3 (63.18%), mainly due to the larger amounts of satellites (over 38 Mb in each T2T genome) captured in the T2T genomes compared to BTx623-v3 (around 19 Mb of satellites). Genes in our T2T genomes were predicted using BRAKER combining evidences from protein homology and RNA-seq data with *ab initio* prediction. A total of 35,695 and 36,950 protein-coding genes were identified in the BTx623-T2T genome and Ji2055-T2T genome, respectively (Table S6). The majority (∼ 83%) of these annotated genes had RNA-seq data support.

### T2T genome identified assembly errors in previous reference genome

Compared to BTx623-v3, the BTx623-T2T genome contained 36.25 Mb of newly assembled sequence. Most (94.10%) of the newly assembled sequence were repeat elements, including retroelements (44.01%) and satellites (45.34%) (Figure 1B, Table S7). These newly assembled sequence was mainly distributed around centromeric regions (82.12%) (Figure 1C). A total of 133 genes were identified in the newly assembled sequence with around 65% of them supported with RNA-seq data. These newly identified genes were predicted to play a role in transmembrane transport, regulation of transcription, developmental process, etc. The BTx623-T2T genome identified the misorientation of four genomics regions around the centromeres of chromosome 1 (7.39 Mb), 5 (20.80 Mb), 6 (6.28 Mb) and 7(13.13 Mb) in BTx623-v3, in addition to mispositioning of two over 1 Mb sequence segments on chromosome 5 and 7, and the absence of hundreds of sequence segments (Figure 1D). Correcting these assembly errors in the reference genome is critical for exploiting genetic information in these complex regions for functional genomics research in sorghum.

The centromere size of BTx623-T2T varied from 2.24 Mb on chromosome 1 to 13.70 Mb on chromosome 4 (Table S5). DNA sequence in centromere was mainly composed of satellite and retrotransposon such as Gypsy and Copia (Table S8). However, the content of these repeat elements differed among chromosomes. Gypsy accounted for more centromeric sequence than satellite did in chromosome 3, 5, 6, 7, 8 and 9, while satellite was the most abundant component of centromeric sequence in chromosome 1, 2, 4 and 10. A total of 134 genes were identified in centromeric regions of BTx623-T2T. These genes were enriched with biological functions such as reproductive process, response to stimuli, developmental processes, etc, suggesting they are critical to fundamental biological processes in sorghum.

### Sequence variation between the sorghum T2T genomes

The assembly of two sorghum T2T genomes allows us to investigate sequence variation across the sorghum genome with a focus on centromeric regions. Substantial sequence variation was observed between the BTx623-T2T genome and the Ji2055-T2T genome. The size of centromeres varied between the two T2T genomes, particularly for chromosome 1, 5, and 7 (Figure 1E). However, the sequence composition of centromeres was largely stable between the corresponding chromosomes of the two genomes (Figure 1F), suggesting the variation of centromere size is unlikely due to expansion of a particular class of repeat element. Most of the genes (84.96%) in centromeres were syntenic between BTx623-T2T and Ji2055-T2T, possibly due to limited recombination in these regions. Sequence comparison of the two T2T genomes identified a total of six large inversions (> 50 Kb) (Figure S7, Table S9). However, none of them overlapped with the centromeric regions.

In summary, this study assembled complete genome sequence of the sorghum reference genome, BTx623 and a popular Chinese inbred line, Ji2055. These two high-quality T2T genomes could serve as the new reference genomes to guide biological discovery and unlock the full potential of global sequence variation for genetic improvement of sorghum.

## AUTHOR CONTRIBUTIONS

Yongfu Tao designed the project. Chuanzheng Wei, Lei Gao, Ruixue Xiao, Bingru Chen, Jihong Li and Yanbo Wang analyzed the sequence. Chuanzheng Wei and Wenhui Zou performed glasshouse experiments. Yongfu Tao, Emma Mace and David Jordan supervised the work. Chuanzheng Wei and Yongfu Tao wrote the manuscript. All authors read and approved the final manuscript.

## ACKNOWLEDGMENTS

The authors thank Prof. Weihua Pan (Agricultural Genomics Institute at Shenzhen, Chinese Academy of Agricultural Sciences) for the discussion on T2T genome assembly. This work was supported by the National Natural Science Fund for Excellent Young Scientists Fund Program (Overseas), the startup package from Agricultural Genomics Institute at Shenzhen, Chinese Academy of Agricultural Sciences and National Natural Science Foundation of China (No.32372176).

## DATA AVAILABILITY STATEMENT

The sequencing data, genome assembly and annotation data generated in this study were deposited in the Genome Warehouse in National Genomics Data Center, Beijing Institute of Genomics, Chinese Academy of Sciences / China National Center for Bioinformation, under accession number PRJCA024204 that is publicly accessible at https://ngdc.cncb.ac.cn/gwh. The codes used in this study can be found at https://github.com/ChuanzhengWei/sorghum_T2T.

## ETHICS STATEMENT

No animals or humans were involved in this study.

## SUPPORTING INFORMATION

The online version contains supplementary figures and tables available.

Figure S1. The seeds of Ji2055 (left) and BTx623 (right).

Figure S2. HiC interaction figure of BTx623 and Ji2055.

Figure S3. Circos plot shows genome feature of BTx623-T2T. (A) Chromosome, (B) Centromere and telomere. (C) Gene density. (D) Density of repeat elements. (E) Density of gyspy. (F) Density of Copia. (G) Density of DNA transposon. (H) GC content.

Figure S4. Mapping rate and coverage rate of 44 sorghum lines against three sorghum genomes.

Figure S5. Distribution of different types of repeat element around centromere regions of BTx-623. Motif includes *PSau3A10* and *pSau3A9*.

Figure S6. Distribution of different types of repeat element around centromere regions of Ji2055 Motif includes *PSau3A10* and *pSau3A9*.

Figure S7. Sequence variation between BTx623-T2T and Ji2055-T2T.

Table S1. Summary of sequencing data generated in this study.

Table S2. Summary statistics of T2T sorghum genome assemblies.

Table S3. Summary of the mapping rate and coverage of 44 sorghum re-sequencing data.

Table S4. Summary of predicted telomeres in our T2T assemblies.

Table S5. Summary of predicted centromeres in our T2T assemblies.

Table S6. Genome annotation of our T2T genomes.

Table S7. Annotation of repeat elements of newly identified sequence in BTx623-T2T.

Table S8. Composition of centromeres in the two sorghum T2T genomes.

Table S9. Sequence variation identified between BTx623-T2T with Ji2055-T2T.

