## Supplementary material for "Complete telomere-to-telomere assemblies of two sorghum genomes to guide biological discovery"

### Supplementary notes

#### Plant material preparation and genome sequencing

The sorghum seeds of BTx623 and Ji2055 were planted and grown in a growth chamber at 25°C with a 16 h light/8 h dark photoperiod setting at Agricultural Genomics Institute at Shenzhen, Chinese Academy of Agricultural Sciences, Shenzhen, Guangdong, China. Fresh young leave at 20 day after emergence were collected and frozen immediately in liquid nitrogen for DNA extraction. High molecular weight (HMW) DNA was isolated from the leaf tissue using a modified cetyltrimethylammonium bromide (CTAB) method .

The extracted HMW DNA was used to construct sequencing libraries for different sequence platforms. SMRTbell libraries of 15-20kb were constructed for PacBio sequencing according to the manufacturer's instructions. The PacBio Revio sequencer was used to sequence the libraries, generating 44.62 Gb and 48.49 Gb HiFi data for BTx623 and Ji2055, respectively. For ONT ultra-long sequencing, sequencing libraries were constructed with HMW DNA, which were then sequenced using Oxford Nanopore Technology GridION X5/PromethION sequencer, producing 74.15 Gb data for BTx623 and 158.69 Gb data for Ji2055. Hi-C sequencing libraries were prepared using a standard protocol [1]. Illumina Novaseq 6000 sequencer was used to sequence the Hi-C libraries, resulting in 37.24Gb and 116.72Gb data for BTx623 and Ji2055, respectively. Illumina sequencing libraries were constructed according to the standard protocol and sequenced with Illumina Novaseq 6000 sequencer, generating paired-end reads. All the sequencing work was conducted at Kindstar Sequenon Biotechnology Co., LTD (Wuhan, China).

#### Assembly of T2T genomes

The ONT ultralong reads were assembled using NextDenovo (v2.5.2, parameters: genome_size=750m, read_cutoff=50k) [2]. The PacBio HiFi reads were assembled using Hifiasm (v0.19.7-r598) and verkko (v1.4.1) with default setting, respectively [3,4]. The primary contig genomes (draft assembly v1) generated by Hifiasm had best quality, and therefore were used as backbone for further assembly analysis. The Hi-C reads were filtered using fastp (v0.23.4) before being utilized to correct, cluster, and orient the v1 contigs by LACHESIS [5]. As a result, ten large contigs corresponding to 10 chromosomes in sorghum were formed for each genome. Jucierbox (v2.20.00) was employed to further correct the congtigs using Hi-C reads [6]. The orientation of the contigs was adjusted according to BTx623-v3 using seqtk (v1.4-r122) [7]. The contigs generated by NextDenovo and verkko were used to fill the sequence gaps and assemble telomere, leading to improved genome assemblies of the two genomes with only four gaps left for each genome (v2).

Seven of the eight remaining gaps were successfully closed by TGS-GapCloser (v1.2.1) using ONT reads [8]. Our attempt to search for ONT read to fill the last gap on chromosome 4 in BTx623 using blastn (version: 2.14.0+ ) failed [9]. However, we found large amount of repeats of "TAC" and a mispositioning of ~16Kb sequence segment compared to BTx623-v3 surrounding the gap, which could cause the difficulty in closing this gap. To resolve this problem, HiFi reads that mapped to the 500 Kb flanking regions of the gap were extracted and assembled using Hifiasm, which closed the last gap and resulted in gapless assembles of both genomes (v3).

HiFi reads were mapped to the assembled genomes to identify high-coverage regions (HCR) and low-coverage regions (LCR) using minimap2 [10]. PanDepth (v2.19, https://github.com/HuiyangYu/PanDepth) was used to calculate coverage depth of HiFi reads based on a sliding window of 10 Kb with a step size of 1 Kb. HCR were defined as regions with coverage of HiFi reads higher than two times of genome-wide average. LCR were defined as regions with coverage of HiFi reads lower than one third of genome-wide average. Two HCR on chromosome 1 and chromosome 9 were identified in the assembled Ji2055 genome, while only one HCR was identified in BTx623, corresponding to the HCR on chromosome 1 in Ji2055. Sequence analysis identified multiple repeat of 45S rDNA and 5S rDNA in the HCR on chromosome 1 and chromosome 9, respectively. The high coverage in the regions could be due to the repetitive nature of the sequence in the regions and bias from the sequence alignment tool. A total of thirteen LCR were found in the two genomes, which were further corrected using HiFi reads and ONT reads (v4). The two v4 genomes were polished with NextPolish (v1.4.1, parameters:-min_read_len 1k -max_depth 100), NextPolish2 (v0.2.0) and pilon (v1.24) using HiFi reads and Illumina short reads to obtain the final T2T assembly of BTx623 and Ji2055 [11,12].

**Quality assessment of the assembled genomes**

Coverage depth was estimated for both HiFi reads and ONT reads. The raw HiFi read and ONT reads were mapped to their corresponding T2T genome using minimap2. Coverage depth of the reads across the genome was calculated with PanDepth based on a sliding window of 10 Kb with a step size of 1 Kb. The base accuracy rate of the genomes was estimated based on a k-mer based approach in merquery (v1.3) using HiFi reads [13].

BUSCO (v5.5.0) was used to assess the completeness of the two T2T genomes using embryophyte_odb1o database [14,15]. LTR_FINDER_parallel (v1.1) was employed to identify LTR in the two genomes, and LAI value was calculated with LTR_retriever (v2.9.5 ) to evaluate the continuity of the genomes [16,17]. The completeness of the T2T genomes was further assessed using the published short reads data of 44 sorghum genomes. These short reads were mapped to our T2T genomes and Btx623-v3 using bwa (v0.7.17-r1188) [18]. Mapping rate and coverage were measured with PanDepth.

**Genome annotation**

To annotate repeat sequence in sorghum genomes, a hybrid approach combining homology search and *de novo* prediction was used. Homology prediction was conducted with RepeatMasker (v4.1.5, http://www.repeatmasker.org) according to the RepBase library. *De novo* prediction based on sequence feature was performed using RepeatModeler (v2.0.5, http://www.repeatmasker.org). Results from the two methods were combined to obtain the final set of repeat sequence. For gene annotation, RepeatMasker (v4.1.5) and RepeatModeler (v2.0.5) were employed to soft mask the genomes. Genes in the softly masked genomes were predicted with BREAKER (v3.0.3) [19]. Gene models of *Zea mays* (AGPv4), *Oryza sativa* (v7.0), *Setaria italica* (v2.2) and *Brachypodium distachyon* (v3.2) were extracted from phytozome (https://phytozome-next.jgi.doe.gov/). Over 600 Gb of RNA-Seq data in sorghum was extracted from National Center for Biotechnology Information (NCBI, https://www.ncbi.nlm.nih.gov/). These gene models and RNA-seq data were used to train AUGUSTUS (http://augustus.gobics.de/) and GeneMark-ETP to perform gene prediction [20]. Coverage rate of RNA-seq reads in predicted genes was summarized using in-house scripts. Genes with more than 50% of predicted mRNA sequence covered by RNA-seq were considered as being supported by RNA-seq data. Gene Ontology (GO) enrichment analysis of genes was conducted using Gene Functional Annotation for Plants (GFAP) [21].

**Identification of centromere and telomere**

Telomeres were identified using quarTeT (v1.1.6, parameters: TeloExplorer -c plant) with command “seqtk telo -m TTTAGGG”. Centromeres were identified with CentroMiner in quarTeT [22]. The borders of centromeres were further refined to include centromere-specific repetitive elements, *PSau3A10* and *pSau3A9.*

**Genome comparison**

Sequence comparison between BTx623-v3 and BTx623-T2T was conducted with minimap2 (parameters: -ax asm5 -t 64 --eqx). Genomic regions with sequence divergence higher than 0.1% was stored in SAM format. Structural variation was identified using SyRI (v1.6.5 ) with default setting, and summarized using in-house scripts [23]. Unaligned regions longer than 1,000 bp in BTx623-T2T were defined as newly assembled sequence. Sequence comparison between Ji2055-T2T and Btx623-T2T was conducted using the same approach. Results of the sequence comparison were visualized using Plotsr [24].

### Supplementary Tables

Table S1. Summary of sequencing data generated in this study.

| Genome | Platform | Reads Number | Base Number | N50 (bp) |
|---|---|---|---|---|
| BTx623 | ultra-long ONT | 858,361 | 79,616,980,308 | 38,434 |
| | Pacbio HiFi | 2,483,114 | 47,914,746,568 | 19,704 |
| | Hi-C | 264,787,247 | 39,982,874,297 | NA |
| | Illumina | 129,964,121 | 19,624,582,271 | NA |
| Ji2055 | ultra-long ONT | 12,437,131 | 170,396,377,137 | 100,001 |
| | Pacbio HiFi | 2,743,084 | 52,065,988,258 | 19,383 |
| | Hi-C | 829,987,574 | 125,328,123,674 | NA |
| | Illumina | 124,875,982 | 18,856,273,282 | NA |

Platform indicates sequencing platforms used to generate sequencing data.

Table S2. Summary statistics of T2T sorghum genome assemblies.

| Parameters | BTx623-T2T | Ji2055-T2T |
|---|---|---|
| Chromosomal genome size (bp) | 719,899,664 | 722,964,161 |
| Contig N50 (bp) | 72,850,042 | 73,908,011 |
| Contig L50 | 5 | 5 |
| LTR Assembly Index | 25.17 | 24.07 |
| HiFi mapping rate | 100% | 100% |
| ONT mapping rate | 99.99% | 99.85% |
| BUSCO | 98.50% | 98.60% |
| Genome quality value score | 70.93 | 71.98 |
| Base error rate | $3.87\times10^{-7}$ | $3.99\times10^{-7}$ |

N50 means the sequence length of the shortest contig at 50% of the total assembly length. L50 means the least number of contigs, of which the length sum accounts for 50% of genome size. Genome quality value score and base error rate of our T2T genomes were estimated using corresponding HiFi reads.

Table S3. Summary of the mapping rate and coverage of 44 sorghum re-sequencing data.

| Sample ID | Mapping rate (%) | | | Coverage (%) | | |
|---|---|---|---|---|---|---|
| | BTx623-v3 | BTx623-T2T | Ji2055-T2T | BTx623-v3 | BTx623-T2T | Ji2055-T2T |
| SRR998970 | 97.41 | 98.96 | 99.05 | 90.49 | 94.68 | 94.81 |
| SRR998971 | 97.62 | 99.35 | 99.36 | 88.19 | 92.08 | 91.97 |
| SRR998972 | 97.63 | 99.49 | 99.46 | 89.92 | 93.98 | 93.47 |
| SRR998984 | 95.67 | 99.44 | 99.42 | 73.6 | 76.46 | 76.71 |
| SRR998985 | 96.81 | 99.53 | 99.57 | 90.16 | 95.21 | 94 |
| SRR998993 | 97.58 | 99.27 | 99.34 | 66.46 | 85.13 | 90.74 |
| SRR998994 | 97.59 | 99.44 | 99.43 | 85.22 | 93.09 | 92.95 |
| SRR998995 | 97.96 | 99.64 | 99.56 | 87.91 | 91.92 | 91.27 |
| SRR998996 | 97.33 | 99.20 | 99.26 | 86.59 | 90.35 | 90.32 |
| SRR998997 | 97.90 | 99.49 | 99.55 | 88.3 | 92.37 | 93.03 |
| SRR998998 | 98.19 | 99.56 | 99.49 | 88.48 | 92.48 | 92.12 |
| SRR998999 | 97.46 | 99.04 | 99.04 | 85.7 | 89.51 | 88.93 |
| SRR999000 | 98.00 | 99.55 | 99.46 | 89.47 | 93.54 | 92.7 |
| SRR999001 | 97.92 | 99.38 | 99.35 | 87.61 | 91.53 | 90.92 |
| SRR999002 | 97.69 | 99.46 | 99.37 | 87.33 | 91.21 | 90.36 |
| SRR999003 | 97.49 | 99.47 | 99.46 | 80 | 83.79 | 83.53 |
| SRR999004 | 97.70 | 99.28 | 99.26 | 88.1 | 92.09 | 91.84 |
| SRR999005 | 97.87 | 99.51 | 99.58 | 88.17 | 92.32 | 91.6 |
| SRR999006 | 97.90 | 99.57 | 99.58 | 87.04 | 91.11 | 90.54 |
| SRR999007 | 98.25 | 99.75 | 99.62 | 88.74 | 93.02 | 91.13 |
| SRR999008 | 97.23 | 98.98 | 98.97 | 84.07 | 87.62 | 87.5 |
| SRR999009 | 97.80 | 99.49 | 99.47 | 88.98 | 93.1 | 91.97 |
| SRR999010 | 98.04 | 99.46 | 99.40 | 87.91 | 91.86 | 91.45 |
| SRR999011 | 97.96 | 99.52 | 99.53 | 89.96 | 94.01 | 93.53 |
| SRR999012 | 97.79 | 99.52 | 99.46 | 89.42 | 93.54 | 92.69 |
| SRR999013 | 98.12 | 99.56 | 99.49 | 89.93 | 94 | 93.06 |
| SRR999014 | 96.66 | 98.19 | 98.18 | 86.89 | 90.72 | 90.56 |
| SRR999015 | 98.16 | 99.56 | 99.57 | 87.8 | 91.93 | 92.5 |
| SRR999016 | 98.02 | 99.45 | 99.33 | 90.35 | 94.4 | 93.55 |
| SRR999017 | 97.74 | 99.55 | 99.54 | 90.17 | 94.2 | 94.39 |
| SRR999018 | 97.39 | 99.39 | 99.38 | 86.6 | 90.48 | 90.42 |
| SRR999019 | 97.87 | 99.43 | 99.45 | 88.55 | 92.68 | 92.54 |
| SRR999020 | 97.82 | 99.40 | 99.38 | 88.3 | 92.26 | 91.7 |
| SRR999021 | 97.09 | 99.07 | 99.05 | 84.19 | 87.8 | 87.82 |
| SRR999022 | 97.99 | 99.56 | 99.47 | 88.95 | 93.05 | 92.04 |
| SRR999023 | 97.32 | 98.83 | 98.82 | 86.35 | 90.02 | 89.94 |
| SRR999024 | 97.76 | 99.30 | 99.29 | 87.51 | 91.39 | 91.16 |
| SRR999025 | 98.04 | 99.41 | 99.43 | 88.11 | 91.97 | 91.64 |
| SRR999026 | 96.32 | 97.80 | 97.79 | 81.93 | 85.28 | 85.15 |
| SRR999027 | 95.09 | 96.88 | 96.83 | 74.59 | 77.33 | 77.42 |
| SRR999028 | 95.47 | 96.75 | 96.70 | 77.23 | 80.3 | 80.38 |
| SRR999029 | 96.92 | 99.31 | 99.42 | 87.83 | 91.74 | 91.71 |
| SRR999030 | 96.41 | 99.32 | 99.28 | 76.27 | 79.72 | 79.87 |
| SRR999031 | 96.99 | 99.28 | 99.25 | 87.21 | 91.02 | 90.9 |
| Average | 97.45 | 99.21 | 99.20 | 86.06 | 90.37 | 90.16 |

Table S4. Summary of predicted telomeres in our T2T assemblies.

| Genome | Chr | Start | End | Number | Start | End | Number |
|---|---|---|---|---|---|---|---|
| BTx623 | Chr01 | 0 | 14,231 | 1,943 | 86,296,768 | 86,304,017 | 1,018 |
| | Chr02 | 0 | 12,215 | 1,626 | 79,321,513 | 79,329,647 | 1,099 |
| | Chr03 | 0 | 10,182 | 1,375 | 80,849,411 | 80,862,180 | 1,683 |
| | Chr04 | 0 | 11,175 | 1,521 | 72,845,387 | 72,850,042 | 584 |
| | Chr05 | 0 | 14,061 | 1,929 | 77,061,176 | 77,069,631 | 1,163 |
| | Chr06 | 0 | 8,502 | 1,147 | 62,672,900 | 62,684,188 | 1,598 |
| | Chr07 | 0 | 7,481 | 1,055 | 69,765,271 | 69,780,702 | 2,113 |
| | Chr08 | 0 | 11,885 | 1,615 | 65,661,732 | 65,670,982 | 1,275 |
| | Chr09 | 0 | 6,947 | 939 | 62,453,443 | 62,463,383 | 1,311 |
| | Chr10 | 0 | 4,943 | 627 | 62,871,900 | 62,884,892 | 1,786 |
| Ji2055 | Chr01 | 0 | 6,581 | 948 | 86,898,069 | 86,905,694 | 1,023 |
| | Chr02 | 0 | 8,234 | 1,094 | 79,768,596 | 79,778,901 | 1,418 |
| | Chr03 | 0 | 7,751 | 1,082 | 80,225,284 | 80,234,350 | 1,228 |
| | Chr04 | 0 | 8,439 | 1,167 | 73,898,595 | 73,908,011 | 1,216 |
| | Chr05 | 0 | 6,329 | 839 | 78,369,653 | 78,376,937 | 986 |
| | Chr06 | 0 | 8,036 | 1,117 | 62,787,015 | 62,795,522 | 1,202 |
| | Chr07 | 0 | 8,439 | 1,184 | 68,571,321 | 68,578,541 | 1,016 |
| | Chr08 | 0 | 5,783 | 806 | 65,893,977 | 65,895,442 | 212 |
| | Chr09 | 0 | 7,111 | 957 | 61,573,613 | 61,581,816 | 1,087 |
| | Chr10 | 0 | 9,838 | 1,319 | 64,897,770 | 64,908,947 | 1,506 |

Chr indicates chromosome. Number summarize the number of "CCCTAAA/TTTAGGG" repeats identified in the region.

Table S5. Summary of predicted centromeres in our T2T assemblies.

| Genome | Chr | Start | End | Length | L/S ratio |
|---|---|---|---|---|---|
| BTx623 | Chr01 | 35,483,416 | 39,769,346 | 4,285,930 | 1.31 |
| | Chr02 | 27,699,249 | 39,674,270 | 11,975,021 | 1.43 |
| | Chr03 | 32,260,094 | 42,906,613 | 10,646,519 | 1.18 |
| | Chr04 | 25,174,651 | 38,870,047 | 13,695,396 | 1.35 |
| | Chr05 | 31,373,897 | 40,943,981 | 9,570,084 | 1.15 |
| | Chr06 | 19,686,946 | 25,481,319 | 5,794,373 | 1.89 |
| | Chr07 | 31,309,234 | 38,582,095 | 7,272,861 | 1.00 |
| | Chr08 | 26,227,639 | 33,879,377 | 7,651,738 | 1.21 |
| | Chr09 | 25,290,887 | 34,119,742 | 8,828,855 | 1.12 |
| | Chr10 | 28,955,725 | 34,981,699 | 6,025,974 | 1.04 |
| Ji2055 | Chr01 | 34,335,322 | 40,798,646 | 6,463,324 | 1.34 |
| | Chr02 | 28,099,365 | 40,070,156 | 11,970,791 | 1.41 |
| | Chr03 | 32,045,643 | 42,756,610 | 10,710,967 | 1.17 |
| | Chr04 | 25,822,354 | 39,548,755 | 13,726,401 | 1.33 |
| | Chr05 | 32,483,158 | 41,435,861 | 8,952,703 | 1.14 |
| | Chr06 | 19,565,240 | 25,374,496 | 5,809,256 | 1.91 |
| | Chr07 | 30,004,644 | 36,674,280 | 6,669,636 | 1.06 |
| | Chr08 | 26,272,636 | 33,917,680 | 7,645,044 | 1.22 |
| | Chr09 | 24,792,514 | 33,288,868 | 8,496,354 | 1.14 |
| | Chr10 | 29,268,969 | 36,090,133 | 6,821,164 | 1.02 |

Chr indicate chromosome. L/S ratio was calculated as the ratio between the length of long arm and the length of short arm in the chromosome.

Tables S6. Genome annotation of our T2T genomes.

| Genome annotation | BTx623-T2T | Ji2055-T2T | BTx623-v3 |
|---|---|---|---|
| Total repeat sequence | 66.50% | 65.22% | 63.18% |
| Retroelements | 50.16% | 49.10% | 50.20% |
| DNA transposons | 9.42% | 9.15% | 9.26% |
| Rolling-circles | 0.40% | 0.39% | 0.41% |
| Small RNA | 0.57% | 0.63% | 0.12% |
| Satellites | 5.25% | 5.25% | 2.59% |
| Simple repeats | 0.70% | 0.69% | 0.61% |
| Predicted genes | 35,696 | 36,950 | 34,211 |
| Newly anchored genes | 84 | 56 | NA |
| Newly identified genes | 133 | 1058 | NA |

Table S7. Annotation of repeat elements of newly identified sequence in BTx623-T2T.

| Repeat element | Number | Length | Percentage |
|---|---|---|---|
| Retroelements | 5177 | 16132155 | 44.01% |
| DNA transposons | 688 | 588405 | 1.61% |
| Rolling-circles | 60 | 16671 | 0.05% |
| Small RNA | 330 | 832566 | 2.27% |
| Satellites | 1167 | 16620414 | 45.34% |
| Simple repeats | 564 | 302398 | 0.82% |
| Low complexity | 55 | 2764 | 0.01% |

Percentage means the percentage of genome sequence the repeat element accounts for.

Table S8. Composition of centromeres in the two sorghum T2T genomes.

| Genome | Chr | Gypsy | Satellite | Copia | CMC-EnSpm | Other |
|---|---|---|---|---|---|---|
| BTx623-T2T | Chr01 | 0.53 | 0.41 | 0.05 | 0.00 | 0.01 |
| | Chr02 | 0.27 | 0.58 | 0.11 | 0.04 | 0.01 |
| | Chr03 | 0.46 | 0.42 | 0.07 | 0.04 | 0.00 |
| | Chr04 | 0.35 | 0.56 | 0.06 | 0.02 | 0.01 |
| | Chr05 | 0.43 | 0.49 | 0.07 | 0.01 | 0.01 |
| | Chr06 | 0.53 | 0.37 | 0.10 | 0.00 | 0.00 |
| | Chr07 | 0.52 | 0.43 | 0.04 | 0.00 | 0.01 |
| | Chr08 | 0.54 | 0.41 | 0.04 | 0.01 | 0.00 |
| | Chr09 | 0.45 | 0.48 | 0.04 | 0.03 | 0.01 |
| | Chr10 | 0.61 | 0.33 | 0.05 | 0.00 | 0.00 |
| Ji2055-T2T | Chr01 | 0.47 | 0.47 | 0.05 | 0.00 | 0.01 |
| | Chr02 | 0.27 | 0.58 | 0.11 | 0.04 | 0.01 |
| | Chr03 | 0.47 | 0.42 | 0.07 | 0.04 | 0.00 |
| | Chr04 | 0.35 | 0.56 | 0.06 | 0.02 | 0.01 |
| | Chr05 | 0.47 | 0.45 | 0.06 | 0.01 | 0.01 |
| | Chr06 | 0.53 | 0.37 | 0.10 | 0.00 | 0.00 |
| | Chr07 | 0.52 | 0.42 | 0.05 | 0.00 | 0.00 |
| | Chr08 | 0.54 | 0.41 | 0.04 | 0.01 | 0.00 |
| | Chr09 | 0.47 | 0.46 | 0.04 | 0.02 | 0.01 |
| | Chr10 | 0.62 | 0.32 | 0.05 | 0.01 | 0.01 |

Chr indicates chromosome.

Table S9. Sequence variation identified between BTx623-T2T with Ji2055-T2T.

| Variation type | Count | Length in BTx623-T2T | Length in Ji2055-T2T |
|---|---|---|---|
| Inversions | 52 | 3,000,005 | 2,688,920 |
| Translocations | 68 | 401,210 | 386,418 |
| BTx623_T2T not aligned | 242 | 36,399,047 | / |
| J2055_T2T not aligned | 375 | / | 39,205,336 |
| SNPs | 514,762 | 514,762 | 514,762 |
| Indels | 121,640 | 5,514,545 | 5,110,299 |

**Supplementary Figure**

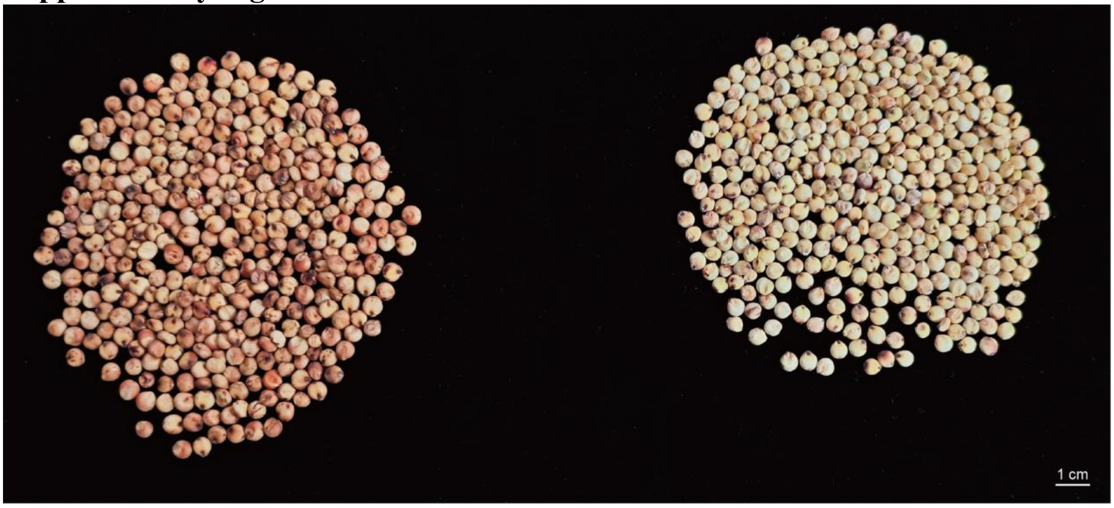


Figure S1 The seeds of Ji2055(left) and BTx623(right).

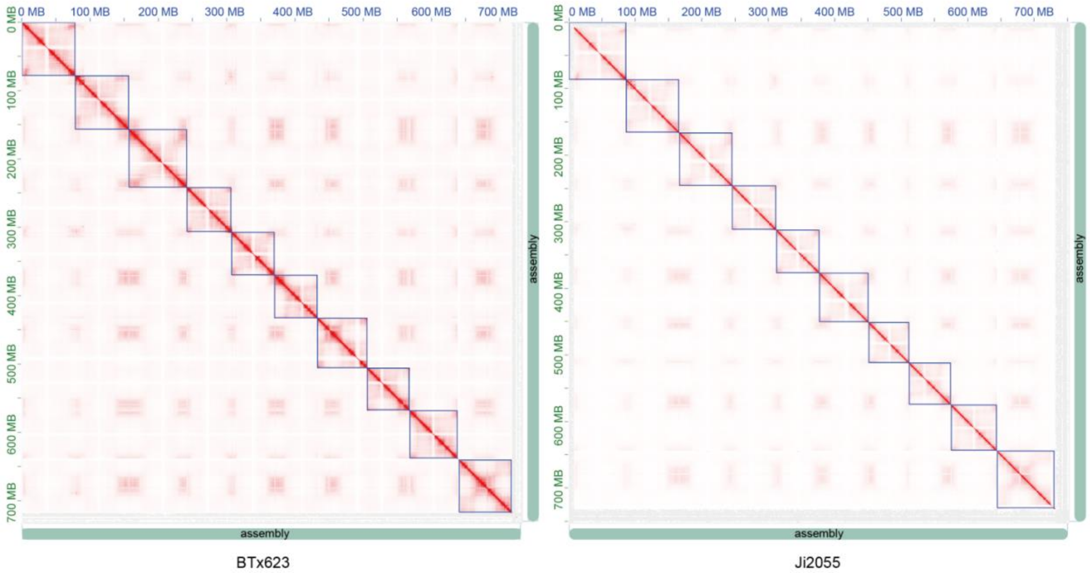


Figure S2 HiC interaction figure of BTx623 and Ji2055.

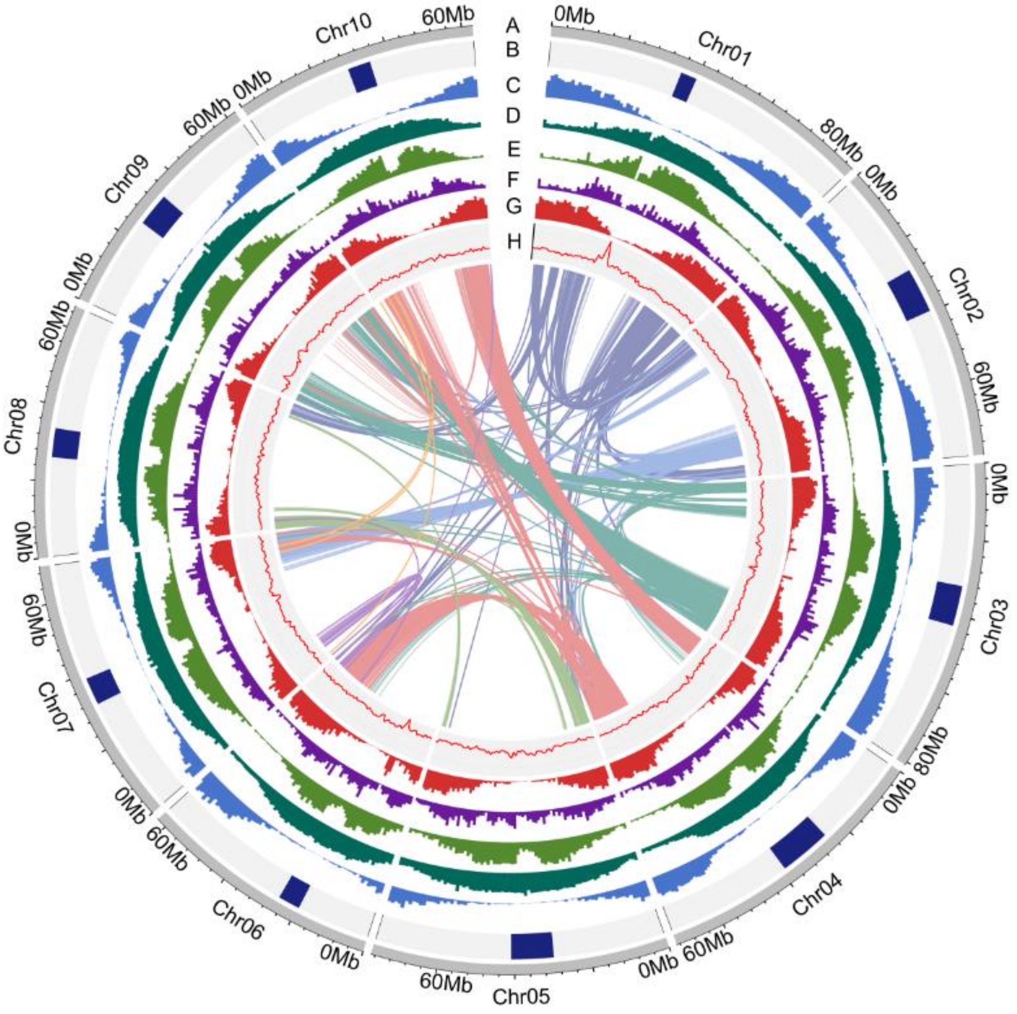


Figure S3 Circos plot shows genome feature of BTx623-T2T. (A) Chromosome, (B) Centromere and telomere. (C) Gene density. (D) Density of repeat elements. (E) Density of gyspy. (F) Density of Copia. (G) Density of DNA transposon. (H) GC content.

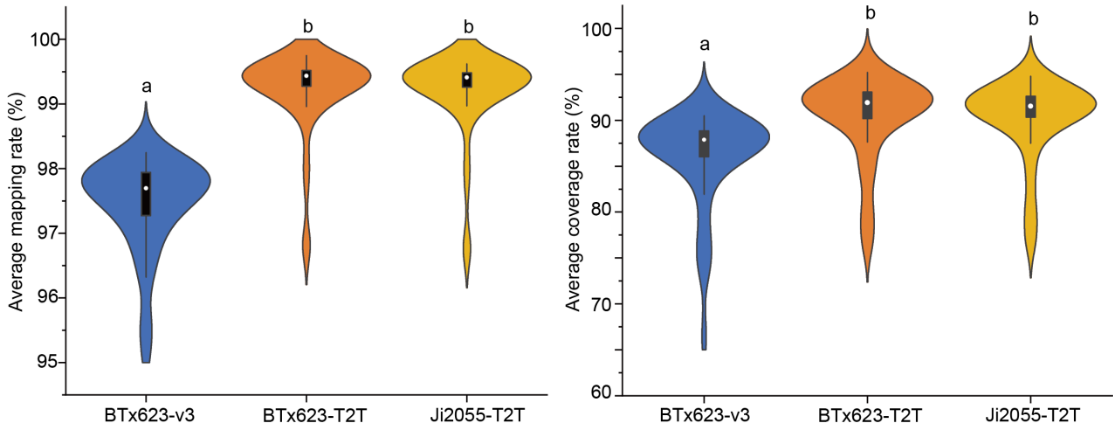


Figure S4 Mapping rate and coverage rate of 44 sorghum lines against three sorghum genomes. Different letters indicate significant difference according to one-way ANOVA analysis followed by Tukey's post hoc test .

263

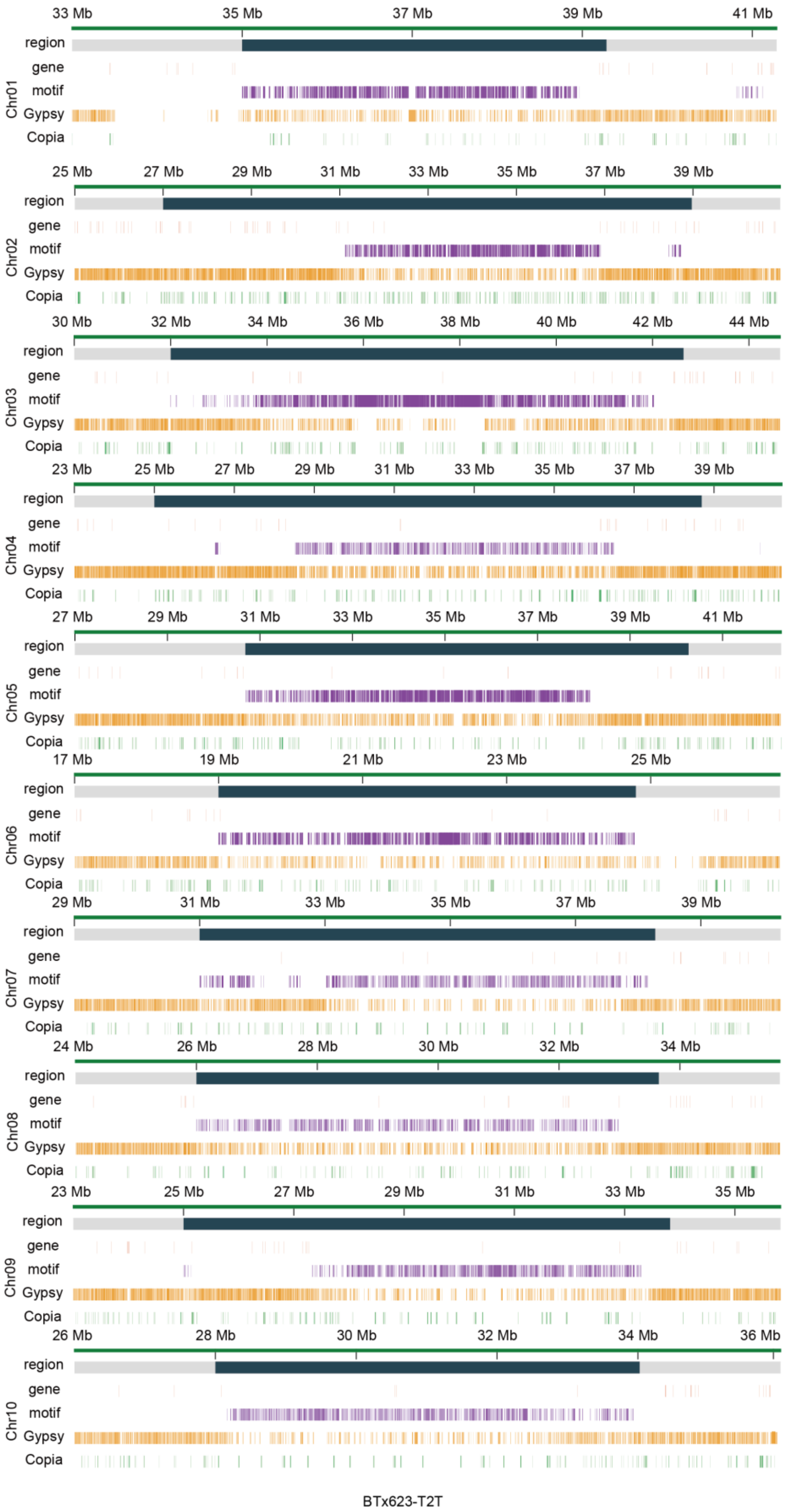


Figure S5 Distribution of different types of repeat element around centromere regions of BTx-623. Motif includes *PSau3A10* and *pSau3A9*.

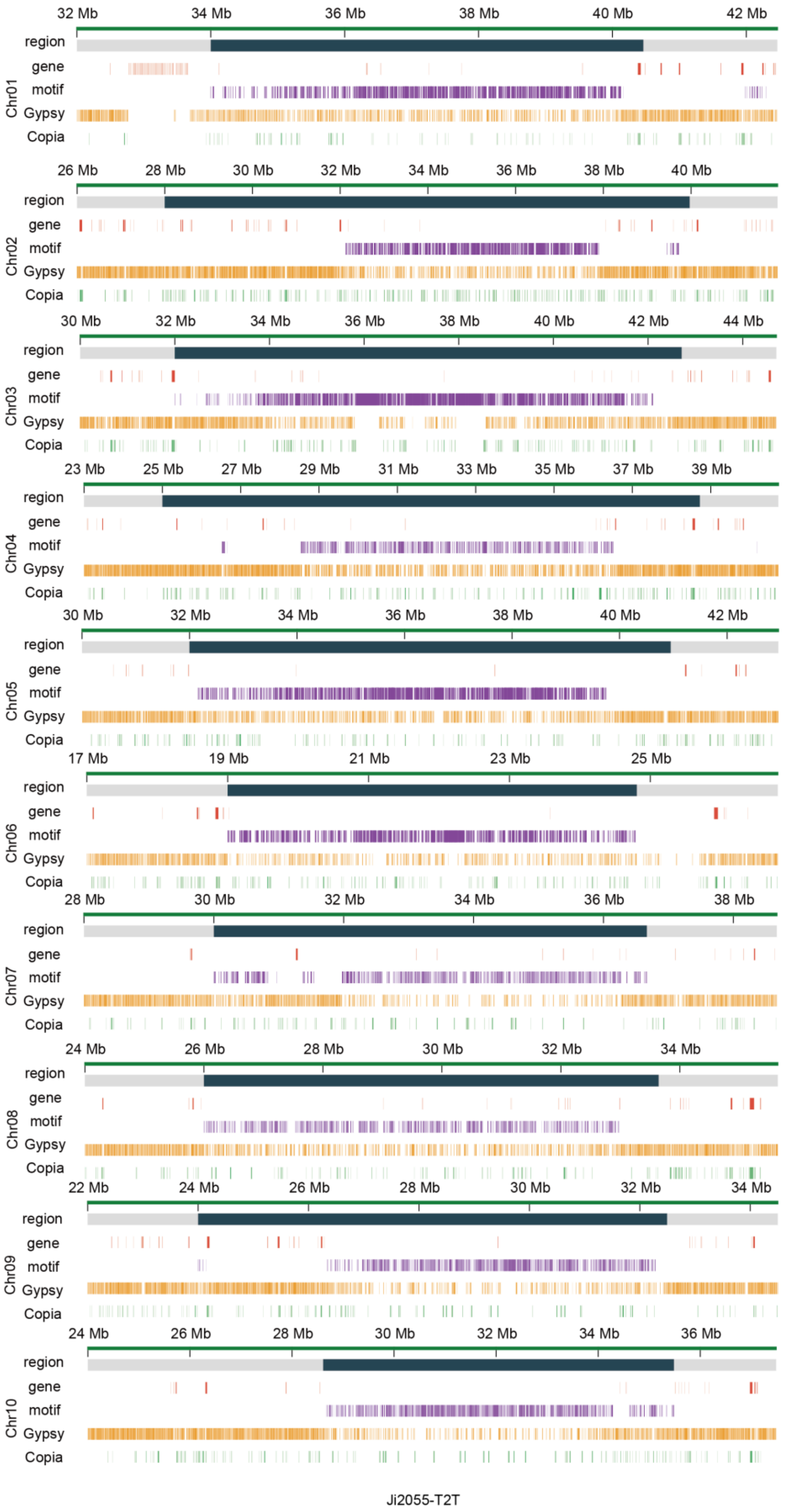


Figure S6 Distribution of different types of repeat element around centromere regions of Ji2055. Motif includes *PSau3A10* and *pSau3A9*.

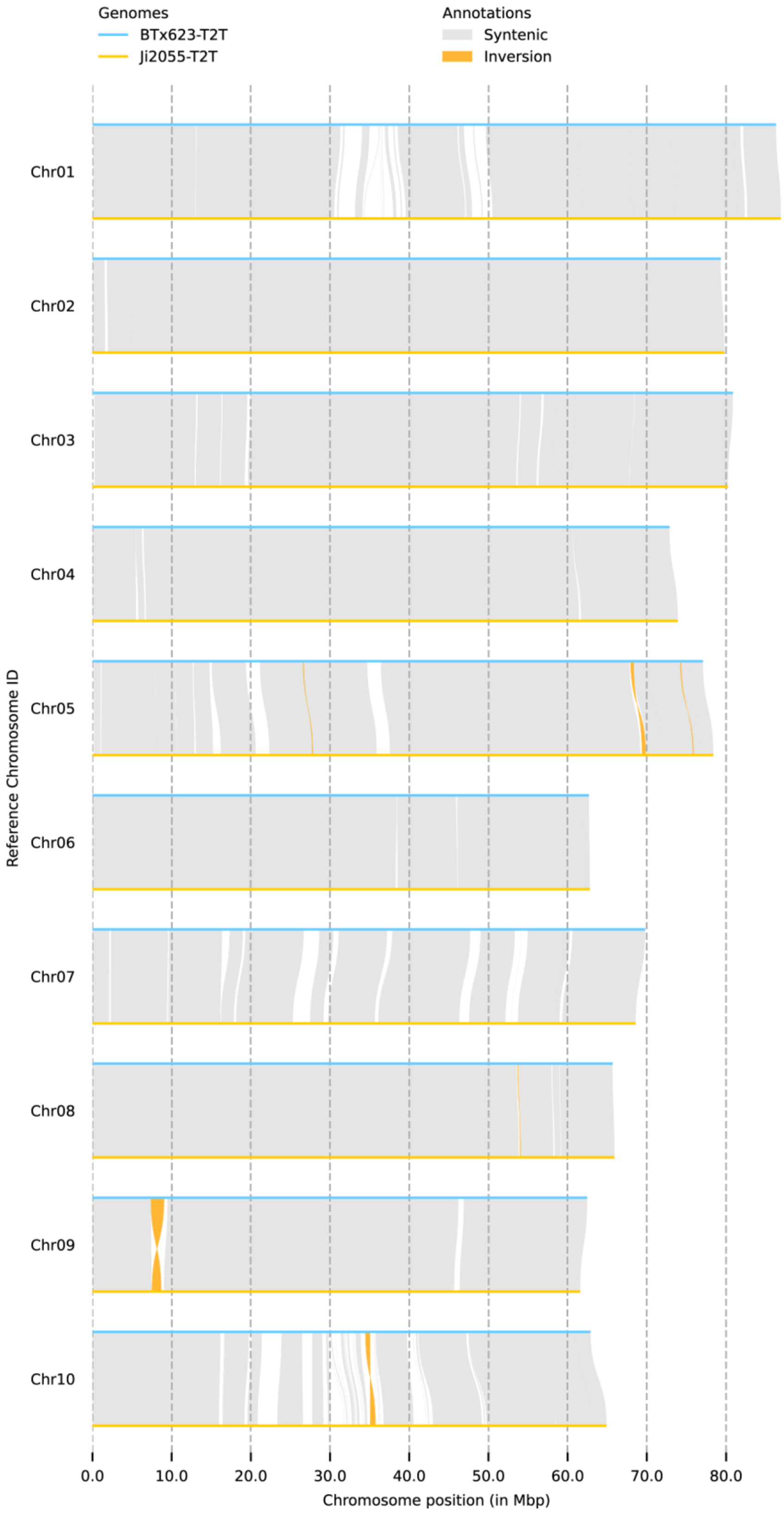


268 Figure S7. Sequence variation between BTx623-T2T and Ji2055-T2T.
